# Guanosine inhibits hepatitis C virus replication and increases indel frequencies, associated with altered intracellular nucleotide pools

**DOI:** 10.1101/2020.02.21.959536

**Authors:** Rosario Sabariegos, Ana M. Ortega-Prieto, Luis Díaz-Martínez, Ana Grande-Pérez, Isabel Gallego, Ana I. de Ávila, María Eugenia Soria, Pablo Gastaminza, Esteban Domingo, Celia Perales, Antonio Mas

## Abstract

In the course of experiments aimed at deciphering the inhibition mechanism of mycophenolic acid and ribavirin in hepatitis C virus (HCV) infection, we observed an inhibitory effect of the nucleoside guanosine (Gua). Here, we report that Gua and not the other standard nucleosides inhibits HCV replication in human hepatoma cells. Gua did not directly inhibit the *in vitro* polymerase activity of NS5B, but it modified the intracellular levels of nucleoside di- and tri-phosphate (NDPs and NTPs), leading to deficient HCV RNA replication and reduction of infectious progeny virus production. Changes in the concentrations of NTP or NDP modified NS5B RNA polymerase activity *in vitro*, in particular *de novo* RNA synthesis and template switching. Furthermore, the Gua-mediated changes were associated with a significant increase in the number of indels in viral RNA, which may account for the reduction of the specific infectivity of the viral progeny, suggesting the presence of defective genomes. Thus, a proper NTP:NDP balance appears to be critical to ensure HCV polymerase fidelity and minimal production of defective genomes.

**Author summary:** Ribonucleoside metabolism is essential for replication of RNA viruses. In this article we describe the antiviral activity of the natural ribonucleoside guanosine (Gua). We demonstrate that hepatitis C virus (HCV) replication is inhibited in the presence of increasing concentrations of this ribonucleoside and that this inhibition does not occur as a consequence of a direct inhibition of HCV polymerase. Cells exposed to increasing concentrations of Gua show imbalances in the intracellular concentrations of nucleoside-diphosphates and triphosphates and as the virus is passaged in these cells, it accumulates mutations that reduce its infectivity and decimate its normal spreading capacity.

## Introduction

Positive-sense single-stranded RNA viruses [(+)ssRNA viruses] are the most abundant pathogens for humans. The hepatitis C virus (HCV) is a hepacivirus that belongs to the *Flaviviridae* family of (+)ssRNA viruses. The HCV genome encodes information for the synthesis of ten proteins: core (C), envelope glycoproteins (E1 and E2), an ion channel (p7), NS2 protease, protease/helicase NS3 (and its cofactor NS4A), membrane-associated protein NS4B, regulator of viral replication NS5A, and RNA-dependent RNA-polymerase NS5B [1].

There are several ways to approach the control of RNA viral diseases. Inhibition of HCV functions by direct-acting antiviral agents (DAAs) has yielded sustained virological responses of about 98% [2, 3]. Thus, HCV infection may be targeted for eradication by the combined use of different DAAs directed to viral proteins. However, access to this treatment is not affordable in countries with high prevalence rates, and an effective prophylactic vaccine is not available, making global HCV eradication difficult. Consequently, treatment with a combination of pegylated interferon-alpha (PEG-IFNα) plus ribavirin (Rib) is still in use in several countries with high prevalence rates of HCV infection [4].

Rib displays several mechanisms of antiviral activity [5], a major one being the inhibition of inosine-5’-monophosphate (IMP) dehydrogenase (IMPDH), which converts IMP to xanthosine monophosphate (XMP) and thus is involved in the *de novo* biosynthesis of GTP [6]. Rib also exerts its antiviral activity through lethal mutagenesis [7–10]. In the course of our experiments on the effect of mycophenolic acid and Rib on HCV clonal population HCV p0 [11] we observed that the presence of guanosine (Gua) during viral replication produced a decrease of up to 100 times in infectious progeny production. Although there are Gua derivatives that have antiviral properties, including Rib itself, natural Gua has never been identified as having antiviral activity [5]. The objective of the present study was to quantify the inhibitory role of Gua on HCV, its specificity, and its mechanism of action. We show that i) Gua inhibits infectious HCV progeny production but does not inhibit directly the HCV polymerase; ii) Gua alters the intracellular pools of di- and triphosphate ribonucleosides (NDP and NTP); iii) the imbalance of the concentrations of NDP and NTP results in the inhibition of HCV polymerase activity *in vitro*, and iv) Gua treatment is associated with an increase of indel frequency in progeny HCV RNA. The results provide evidence of a metabolism-dependent mechanism of generation of defective HCV genomes.

## Results

### Effect of ribonucleosides on HCV replication

Before studying the possible anti-HCV effect of natural nucleosides we determined their cytotoxicity (CC_50_) on Huh-7.5 reporter cells. The cytotoxicity of Gua, adenosine (Ade), cytidine (Cyt) or uridine (Uri) was analyzed in semiconfluent cell monolayers by exposing cells to different nucleoside concentrations (from 0 μM to 800 μM). Cell viability (CC_50_) was monitored after 72 h of treatment (Table 1) as described in Materials and Methods. Only Ade showed a modest cytotoxicity in the range of concentrations tested.

**Table 1.** Effects of nucleosides on cell viability and HCV replication. CC_50_, IC_50_, and therapeutic index (TI, CC_50_/IC_50_) values are shown for Adenosine (Ade), Cytosine (Cyt), Guanosine (Gua), and Uridine (Uri) in Huh-7.5 reporter cells.

| | $CC_{50}$ (μM) | $IC_{50}$ (μM) | TI |
| --- | --- | --- | --- |
| Ade | $641 \pm 40$ | $108 \pm 7$ | 5.9 |
| Cit | > 800 | > 800 | n.d. |
| Gua | > 800 | $164 \pm 2.4$ | $\geq 4.9$ |
| Uri | > 800 | > 800 | n.d. |

To quantify the inhibition of HCV infectious progeny production in the presence of nucleosides (IC_50_), Huh-7.5 reporter cells were infected with HCV p0 at a multiplicity of infection (m.o.i.) of 0.05-0.1 TCID_50_ per cell in the presence of increasing concentrations of the corresponding nucleoside, and infectious progeny production was measured as described in Materials and Methods. A decrease in the production of HCV infectious progeny was observed for Gua and Ade, whereas Cyt and Uri did not show any effect (Table 1). These data yield a therapeutic index (TI), defined as CC_50_/IC_50_, of 5.9 and ≥ 4.9 for Ade and Gua, respectively (Table 1).

To further explore the effect of ribonucleosides on HCV replication, HCV p0 was subjected to 5 serial passages in Huh-7.5 reporter cells, using an initial m.o.i. of 0.05 TCID_50_ per cell, both in the absence and in the presence of ribonucleosides at 500 μM and 800 μM (Fig 1). Results show a consistent decrease in progeny infectivity as a result of Gua treatment (Fig 1A and 1B), but a sustained viral replication in the presence of Ade, Cyt or Uri (Fig 1C, D). In the presence of 500 μM Gua, a decrease in infectivity was detected although only one of the four replicates yielded values below the detection limit (Fig 1A). A sustained drop in HCV infectivity by Gua 800 μM was achieved, which became undetectable between passages 2 and 4 in all replicates (Fig 1B). Therefore, Gua was the only nucleoside that showed antiviral effect without cytotoxicity.

**Figure 1.**
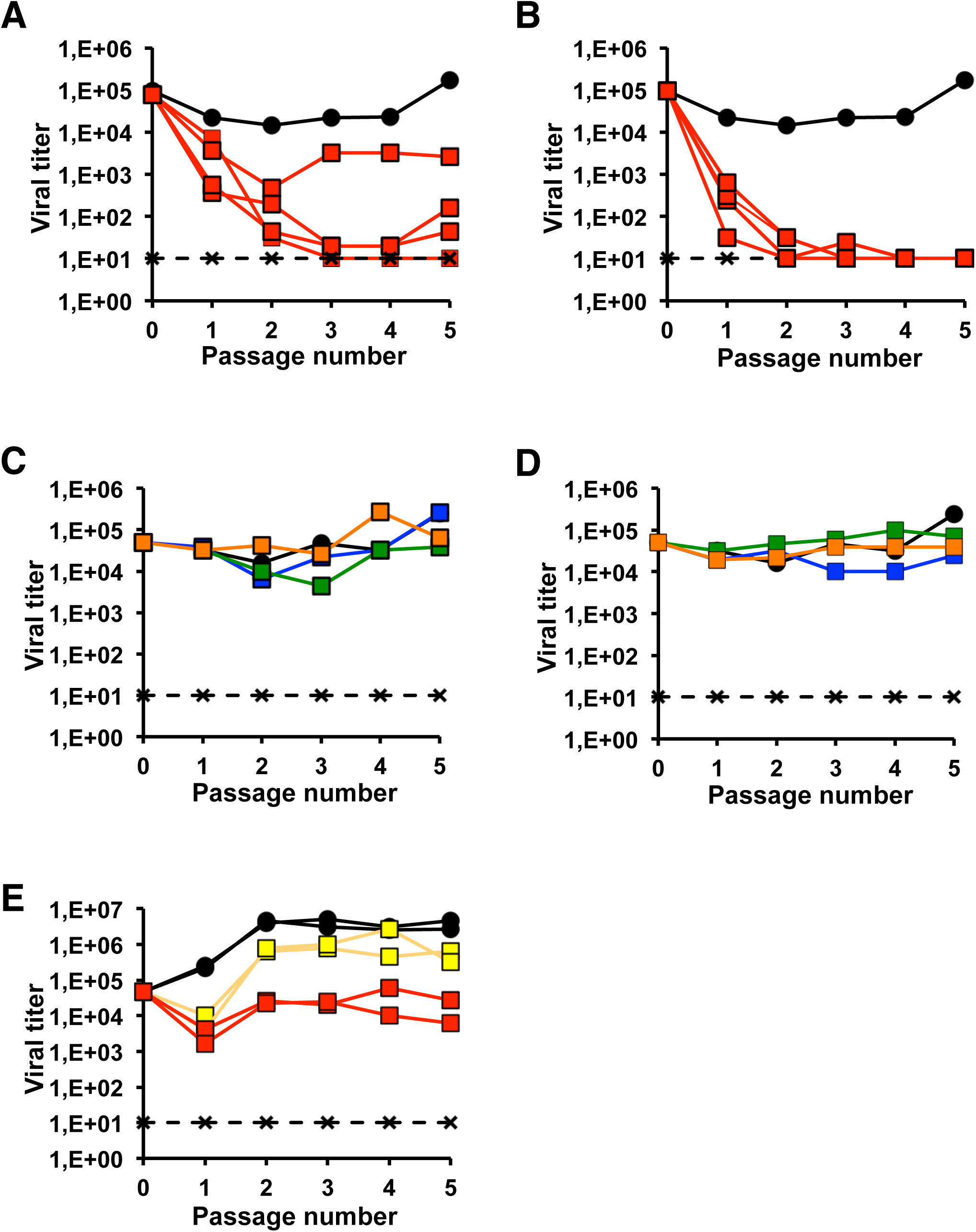
Effect of ribonucleosides on HCV replication. **(A)** and **(B)** Effect of guanosine (Gua) on HCV p0 replication. Infectious progeny obtained in the presence of Gua 500 μM (A) and Gua 800 μM (B); four replicas for each condition are shown (red squares). HCV p0 titer in the absence of treatment (black circles) and values for a HCV lethal mutant GNN (black crosses) are also shown (see Methods). **(C)** and **(D)** Effect of adenosine (Ade, blue squares), cytosine (Cyt, green squares), and uridine (Uri, orange squares) on HCV p0 replication. Infectious progeny obtained in the presence of the corresponding nucleoside at 500 μM (C) and at 800 μM (D). HCV p0 viral titer in the absence of treatment (black circles) and values for a HCV lethal mutant GNN (black crosses) are also shown. **(E)** HCV p100 viral titer in the absence (black circles) or presence of Gua 500 μM (yellow symbols), and Gua 800 μM (red symbols). Two replicates are shown for each condition in presence of nucleosides. The discontinuous horizontal line marks the limit of detection of virus infectivity. Procedures for serial infections and titration of infectivity are detailed in Materials and Methods.

Next, we analyzed the effect of treatment with Gua, Ade, Cyt, and Uri in a surrogate single cycle infection model, taking advantage of spread-deficient bona fide HCV virions bearing a luciferase reporter gene (HCVtcp). This system recapitulates early stages of the infection including viral entry, primary translation and genome replication, overall efficiency of which is proportional to reporter gene activity [12]. The results (Fig 2A) show that a selective HCV entry inhibitor, hydroxyzine, strongly interferes with reporter gene accumulation, as previously documented [13] (Figure 2A). Of the four natural nucleosides, only Gua exerted a significant inhibitory role, as shown by reduced luciferase levels in these cells (Figure 2A), and suggesting that an early step of the infection preceding viral assembly is significantly inhibited by Gua. To further dissect the impact of Gua on HCV replication, we analyzed the effect of Gua, Ade, Cyt and Uri treatment at different times in the replication of a dicistronic subgenomic genotype 2a (JFH-1) replicon bearing a luciferase reporter gene [12]. The objective was to analyze if the effect took place at the level of IRES-dependent translation (5 h post-transfection) or during RNA replication (24 and 48 h post-transfection) [13]. The results (Fig 2B) show that there are no differences among the different treatment points at 5 h post-transfection, which excludes an effect on HCV IRES-dependent RNA translation or any spurious interference with reporter gene expression. However, Gua-treated cells showed a statistically significant 12- and 5-fold reduction in RNA replication at 24 and 48 hours post-transfection respectively (Figure 2B). A modest (±2-fold) but significant reduction was also observed in Ade-treated cells. The fact that Ade treatment does not interfere with HCVtcp (trans-complemented HCV particles) infection suggests that only Gua affects significantly with HCV infection by interfering with viral RNA replication, downstream of viral entry and primary translation.

**Figure 2.**
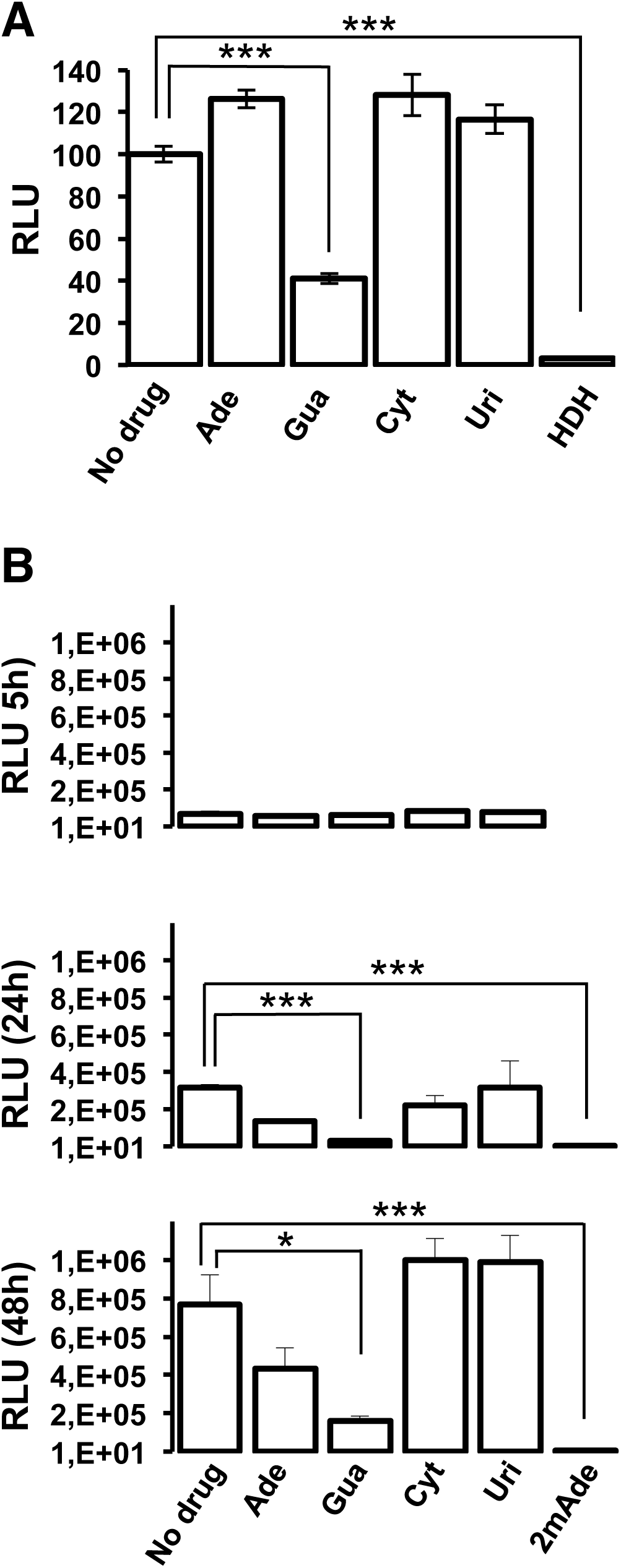
Treatment with Gua causes a reduction in the efficiency of early aspects of the infection. **(A)** Impact of nucleoside treatment on single cycle trans-complemented HCV particles (HCVtcp) infection efficiency. Huh-7 cells were pre-treated with the indicated doses of nucleoside for 20 hours before inoculation with HCVtcp in the presence or absence of the nucleosides. As a positive inhibition control, target cells were treated with the entry inhibitor hydroxyzine (HDH) at the time of infection (5µM). Single cycle infection efficiency was determined by measuring luciferase activity in total cell extracts 48 hours post-inoculation. **(B)** Huh-7 cells were pre-treated with the indicated doses of nucleoside for 20 hours before transfection with in vitro-transcribed subgenomic viral RNA-containing liposomes in the presence or absence of the nucleosides. As a positive inhibition control, target cells were treated with the replication inhibitor 2’-c-methyladenosine (2mAde) (10 µM). Primary translation (5 hours) and RNA replication efficiency (24 and 48 hours) was determined by measuring luciferase activity in total cell extracts at different times post-transfection. Data are shown as average and standard deviation of two experiments performed in triplicate (N=6). Significance (Student’s T-test): ***p<0.0005; *p<0.05.

### Effect of Gua on a high fitness HCV population

The HCV p100 virus [HCV p0 passaged 100 times in Huh-7.5 reporter cells], shows a relative fitness that is 2.2 times higher than that of the HCV p0 parental population [14]. Since viral fitness can influence the response of the virus to antiviral agents [15–17], HCV p100 was used to study the response of a high fitness HCV to Gua. For this, HCV p100 was subjected to 5 serial passages in Huh-7.5 reporter cells using an initial m.o.i. of 0.05 TCID_50_ per cell, both in the absence and in the presence of Gua 500 or 800 μM. The results show a sustained drop in infectivity of 15 and 736 times along the passages as a result of treatment with 500 and 800 μM Gua, respectively. However, no decrease of infectivity below the limit of detection was observed throughout the five passages in any of the replicates (Fig 1E). Only the decrease in progeny production in the presence of 800 µM reached statistical significance (Fig 1E). Thus, the results showed increased resistance of HCV p100 to Gua compared to HCV p0 (compare Fig 1A, 1B, and 1E) as was previously observed with several antiviral drugs [16, 17].

### Effect of Gua on the replication of other RNA viruses

To determine the specificity of the antiviral action exerted by Gua on HCV and to rule out nonspecific effects that could affect any virus, comparative experiments were conducted with foot-and-mouth disease virus (FMDV), lymphocytic choriomeningitis virus (LCMV), and vesicular stomatitis virus (VSV). First, the CC_50_ values of Gua, Ade, Cyt, and Uri were determined for BHK-21 cells, as described in Materials and Methods. The values obtained (Table 2) indicate no detectable cytotoxicity of Gua, Cyt and Uri, and a CC_50_ value of 391 ± 68 μM for Ade. To determine the IC_50_ values of these nucleosides, BHK-21 cells were infected with FMDV, LCMV, and VSV, at an initial m.o.i. of 0.05 TCDI_50_ per cell in the presence of increased nucleoside concentrations and the production of infectious progeny was measured. The values obtained (Table 2) show that all nucleosides lacked inhibitory profile for FMDV. In contrast, purines were inhibitory for VSV, while all nucleosides were inhibitory for LCMV. However, the IC_50_ values were very high and the therapeutic indexes (TI) were consequently low (Table 2).

**Table 2.** Effects of nucleosides on BHK-21 cells viability and VSV, FMDV, and LCMV replication. CC_50_, IC_50_, and therapeutic index (TI, CC_50_/IC_50_) values are shown for Adenosine (Ade), Cytosine (Cit), Guanosine (Gua), and Uridine (Uri) in BHK-21 cells.

|  | CC<sub>50</sub> (μM) | IC<sub>50</sub> (μM) (TI) |  |  |
| --- | --- | --- | --- | --- |
|  |  | VSV | FMDV | LCMV |
| Ade | 391 ± 68 | 650 ± 105.4 (0.6) | > 800 (n.d.) | 213 ± 56.2 (1.8) |
| Cit | > 800 | > 800 (n.d.) | > 800 (n.d.) | 566.4 ± 216.7 (≥ 1.4) |
| Gua | > 800 | 734 ± 59.4 (≤ 1.1) | > 800 (n.d.) | 348 ± 36 (≥ 2.3) |
| Uri | > 800 | > 800 (n.d.) | > 800 (n.d.) | 157.8 ± 7.6 (≥ 5.1) |

n.d. Not Determined.

As an additional control for the specificity of HCV inhibition by Gua, the response of VSV, FMDV and LCMV to nucleoside treatment in serial infections was studied. BHK-21 cells were infected with FMDV, LCMV, and VSV with an initial m.o.i. of 0.05 TCDI_50_ per cell, and were subjected to 3 passages both in the absence and presence of nucleosides at a final concentration of 800 μM. The analysis of the viral populations in passage 3 showed no statistically significant difference from the viral titer obtained in the absence of treatment (Fig 3A-C). Therefore, the results show that the only differences found were those of HCV treatment with Gua at 800 μM (Fig 1A-B). Finally, to rule out that the inhibitory effect of Gua on HCV was solely due to the action of the nucleoside on the human hepatoma cells used in the experiments, we examined the production of VSV viral progeny in Huh-7.5 reporter cells which this virus also productively infects. High viral titers in the presence of 800 μM Gua were obtained for VSV, confirming a lack of antiviral activity of Gua against this virus also in Huh-7.5 reporter cells (Fig 3D).

**Figure 3.**
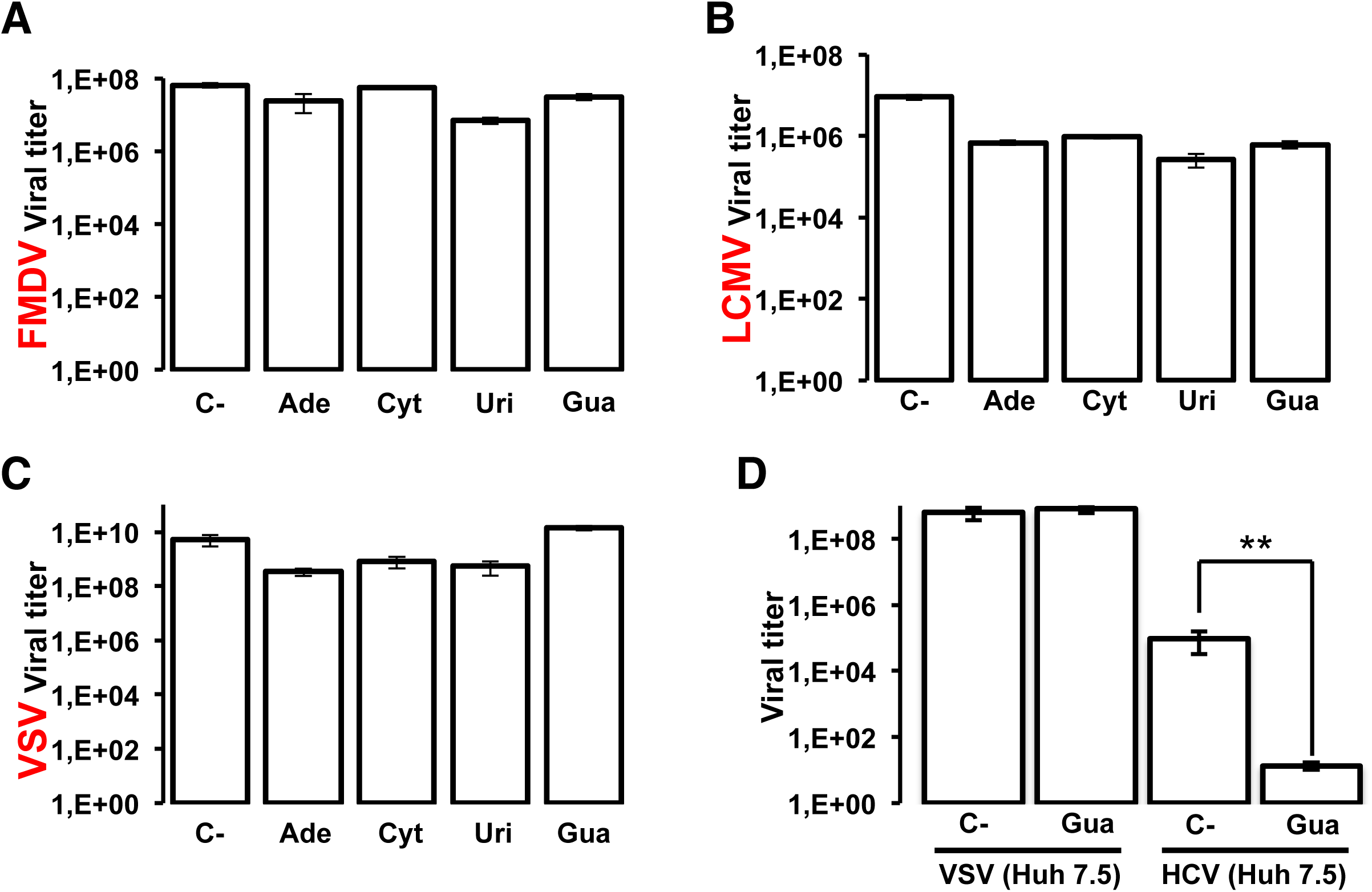
Effect of nucleosides on FMDV, LCMV and VSV replication. Effect of adenosine (Ade), cytidine (Cyt), uridine (Uri), and guanosine (Gua) on the production of infectious progeny of FMDV **(A)**, LCMV **(B)** and VSV **(C)** in BHK-21 cells at an initial m.o.i. of 0.05 TCID_50_ per cell, in the absence (C-) or presence of 800 μM of the indicated nucleoside. Infectivity was determined at passage 3 in the cell culture supernatant as described in Materials and Methods. **(D)** Comparative inhibition of VSV and HCV p0 progeny production in Huh-7.5 reporter cells in the presence of 800 μM Gua. The titer shown for HCV is the average (four replicas) of titers determined at passage 3 in the supernatants of the serial infections (corresponding to Fig 1). Procedures for serial infections and titration of infectivity are detailed in Materials and Methods. Significance (Student’s T-test): ** p < 0.005.

### Effect of guanosine on HCV NS5B activity

To analyze the mechanism by which Gua inhibits HCV replication we tested the effect of increasing Gua concentrations on HCV polymerase activity *in vitro*. A 570 nt RNA fragment corresponding to the E1/E2 region of the HCV genome [18] was replicated by HCV recombinant NS5B**Δ**21 in the presence of ATP, CTP, GTP, and UTP, and at increasing concentrations of Gua (Fig 4A). NS5B polymerase activity increased with Gua concentration up to 500 μM. Even at 1 mM Gua, the RNA polymerase activity was similar to that obtained in absence of Gua. Only at very high Gua concentration (10 mM) the RNA polymerase activity showed a significant reduction (Fig 4A). Similar results were obtained using the 19-mer oligonucleotide LE19 (Fig 4B). Therefore, according to this *in vitro* RNA synthesis assay, the inhibition of HCV progeny production by Gua cannot be attributed to direct inhibition of the HCV RNA polymerase.

**Figure 4.**
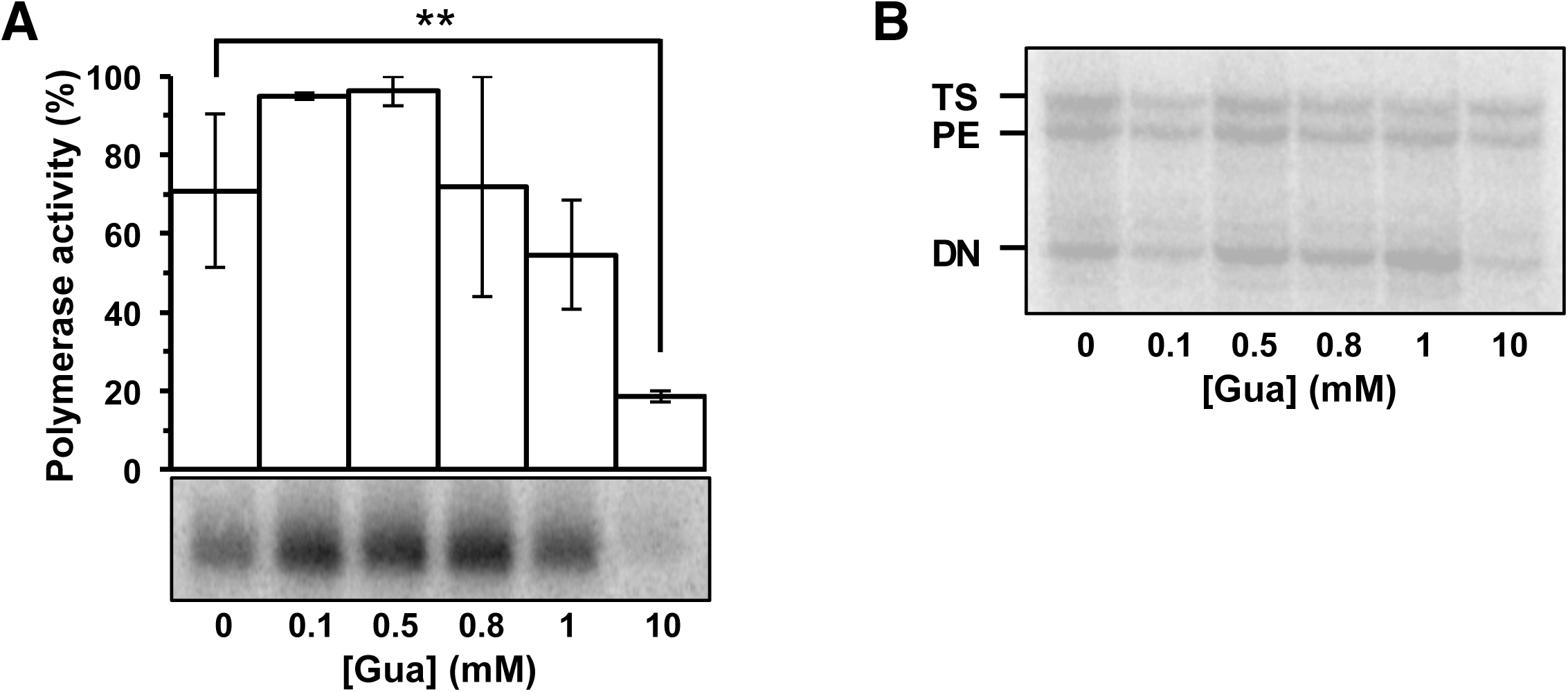
Effect of Gua on NS5BΔ21 RNA polymerase activity. **(A)** Recombinant HCV NS5B**Δ**21 polymerase was added to a reaction containing a 540-nt RNA template [18], the four nucleoside-triphosphates (ATP, CTP, GTP, and UTP) and the indicated concentrations of Gua. Product quantification from three replicates (average ± SEM) and a representative experiment (below) are shown. Polymerase activity is normalized with respect to its maximum activity. The band indicates a new synthesis RNA product of 540 nt length. **(B)** A representative experiment as in A, but using the 19-nt LE19 RNA as a template. DN, PE, and TS indicate reaction products of *de novo* synthesis, primer extension, and template switching, respectively [54]. Procedures are detailed in Materials and Methods. Significance (Student’s T-test): ** p < 0.005.

### Effect of guanosine on intracellular nucleotide pools

To investigate whether HCV replication inhibition by Gua could be related to alterations in di- and triphosphate ribonucleoside intracellular concentrations, the level of NTPs and NDPs in Huh-7.5 reporter cells was determined in the absence of Gua and after 72 h of treatment with 500 or 800 μM Gua. Intracellular nucleoside triphosphate concentrations did not change when the cells were treated with 500 μM Gua. However, 800 μM Gua treatment resulted in a statistically significant 2-fold increase (range from 2- to 2.2-fold) in intracellular concentration of all NTPs (Fig 5A). In the case of NDPs a significant increase of 1.7- to 4.1-fold was observed in cells treated with 500 μM Gua (Fig 5B). Treatment with 800 μM Gua resulted in an increase of the concentration (2.1- to 5.4-fold) of all NDP’s (Fig 5B). Therefore, the presence of Gua in the culture medium increased the intracellular levels of nucleoside di- and tri-phosphates (Supplementary Table). The lowest nucleotide concentration was obtained for CDP and CTP independently of the treatment with Gua.

**Figure 5.**
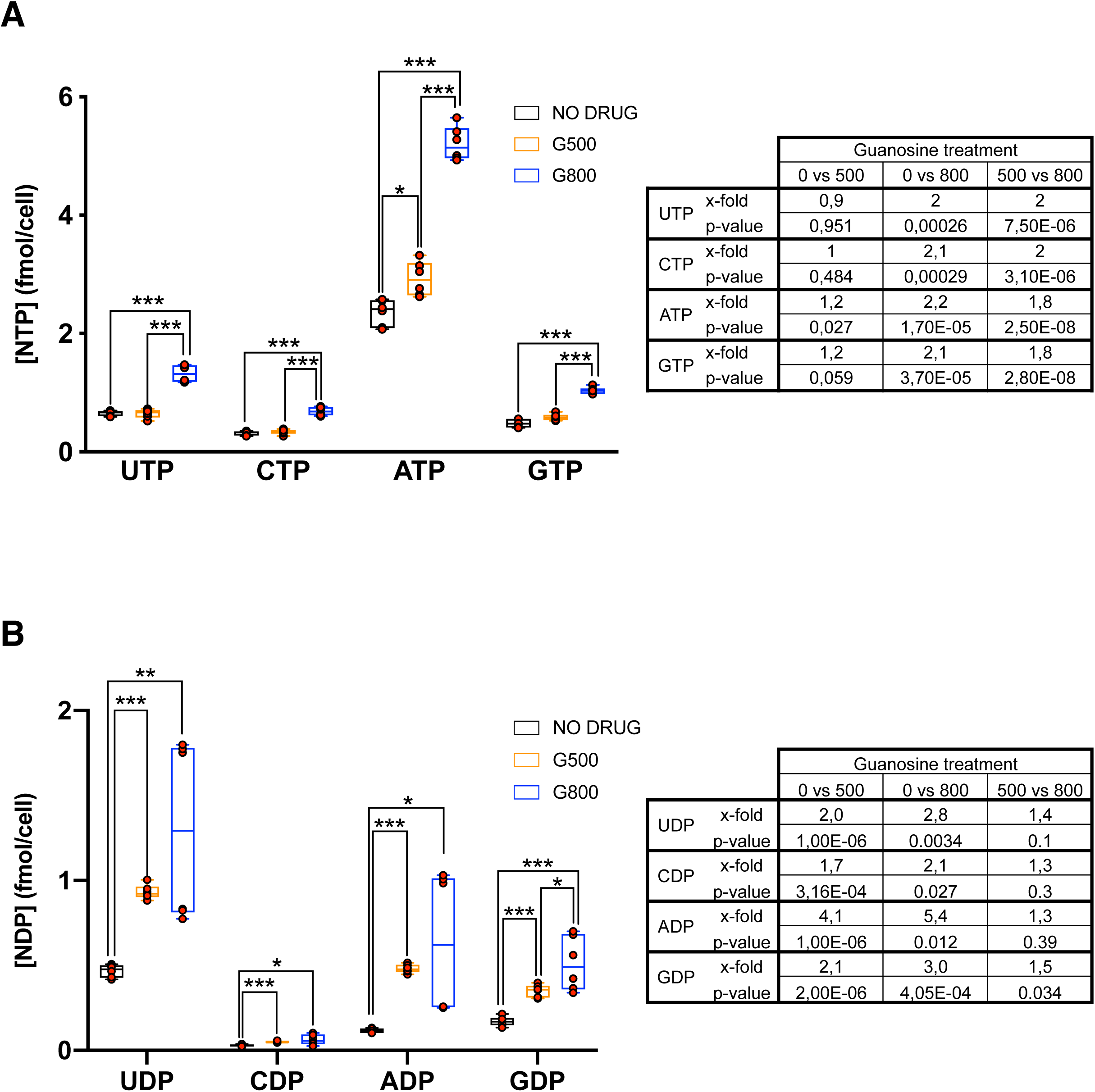
Effect of Gua on intracellular nucleotides. Effect of treatment of Huh-7.5 cells with Gua on the level of intracellular nucleoside-triphosphates (**A**) and intracellular nucleoside-diphosphates (**B**). Data are represented as a box and whisker chart showing distribution of data into quartiles, highlighting the mean and outliers. Error lines indicate variability outside the upper and lower quartiles. Data points (red circles) are grouped in black (no treatment), orange (500 µM Gua) and blue (800 µM Gua) boxes. Significance (Student’s T-test): * p < 0.05; ** p < 0.005; *** p < 0.0005. The table shows the p-value and the fold of difference between treatment conditions for each nucleoside di- and triphosphate.

### Effect of nucleoside di- and triphosphate imbalance on HCV NS5B activity *in vitro*

To explore if changes in nucleotide concentrations might affect HCV polymerase activity, we performed *in vitro* RNA polymerization experiments with recombinant NS5B**Δ**21 in the presence of increasing concentrations of NTPs or NDPs. CTP and CDP were not included in the analyses because they showed small intracellular variations (Supplementary Table) and CTP was chosen as the carrier of the radioisotope. *De novo* (DN), primer extension (PE) and template switching (TS) polymerase activities were measured in the presence of increasing concentrations of the corresponding triphosphate nucleosides (Fig 6). A high UTP concentration of 1 mM slightly but significantly decreased primer extension activity (Fig 6A). However, the main effect of NTP concentration was on the *de novo* RNA synthesis, with a significant decrease at high ATP concentration (Fig 6B), and a significant increase at high GTP concentration (Fig 6C). The increase in the *de novo* RNA synthesis was accompanied by an increase of template switching (Fig 6C).

**Figure 6.**
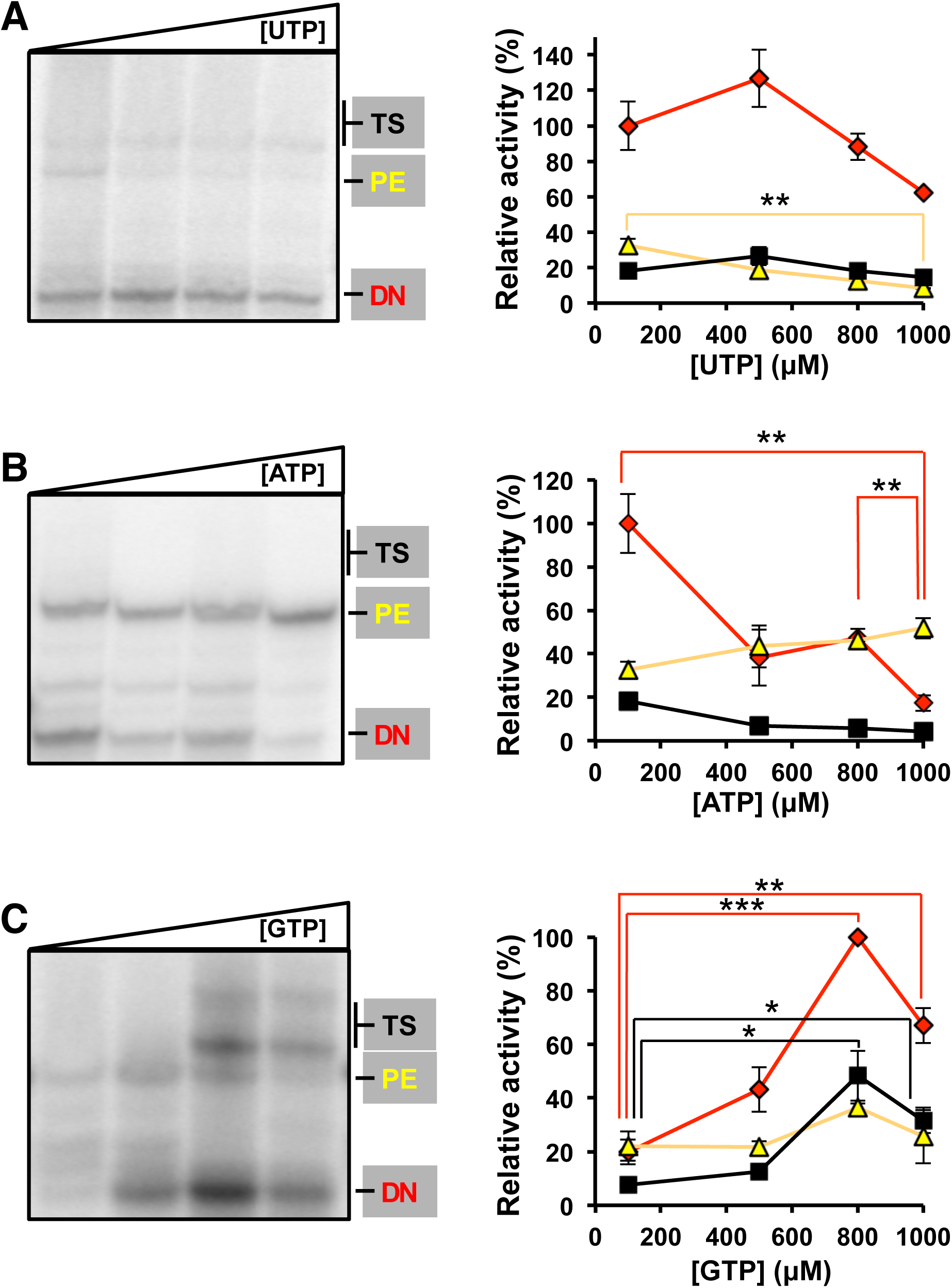
Effect of nucleoside-triphosphate concentration on NS5BΔ21 RNA polymerase activity. (**A)** Polyacrylamide gel showing the products for de novo (DN), primer extension (PE) and template switching (TS) obtained with HCV NS5BΔ21 at increasing concentrations (100, 500, 800, and 1000 μM) of UTP in the presence of radiolabeled α^32^P-CTP. ATP and GTP concentrations were maintained at 100 μM and 500 μM, respectively (left panel). Graphic representation of densitometric values obtained from the electropherogram shown in A (red diamonds, yellow triangles, and black squares correspond to *de novo* (DN), primer extension (PE), and template switching (TS) activities, respectively) (right panel). **(B)** Corresponds to experiments as in **A** but for increasing concentrations of ATP, with UTP and GTP maintained at 100 μM and 500 μM, respectively. **(C)** Corresponds to experiments as in **A** but for increasing concentrations of GTP, with ATP and UTP both maintained at 100 μM. Activities were normalized to their maximum values. Densitometric data represent the mean of at least three independent experiments. Error bars correspond to standard error of the mean. Horizontal lines indicate statistically significant differences (Student’s T-test) between the activity values that link, using the same color code as the activity type. Details of the activity measurements are given in Materials and Methods. Significance (Student’s T-test): *** p < 0.0005, ** p < 0.005, * p < 0.05.

Since increasing concentrations of Gua also altered the intracellular NDP concentrations (Fig 5), we investigated if the presence of increasing concentration of NDPs might affect the NS5B RNA polymerase activity *in vitro*. DN, PE, TS activities were measured in the presence of increasing concentrations of the corresponding diphosphate nucleosides at a fixed nucleoside-triphosphate concentration (Fig 7). The main effect of the presence of NDP was on the *de novo* RNA synthesis, with a significant decrease at high ADP and GDP concentrations. Differences in primer extension and template switching did not reach statistical significance (Fig 7).

**Figure 7.**
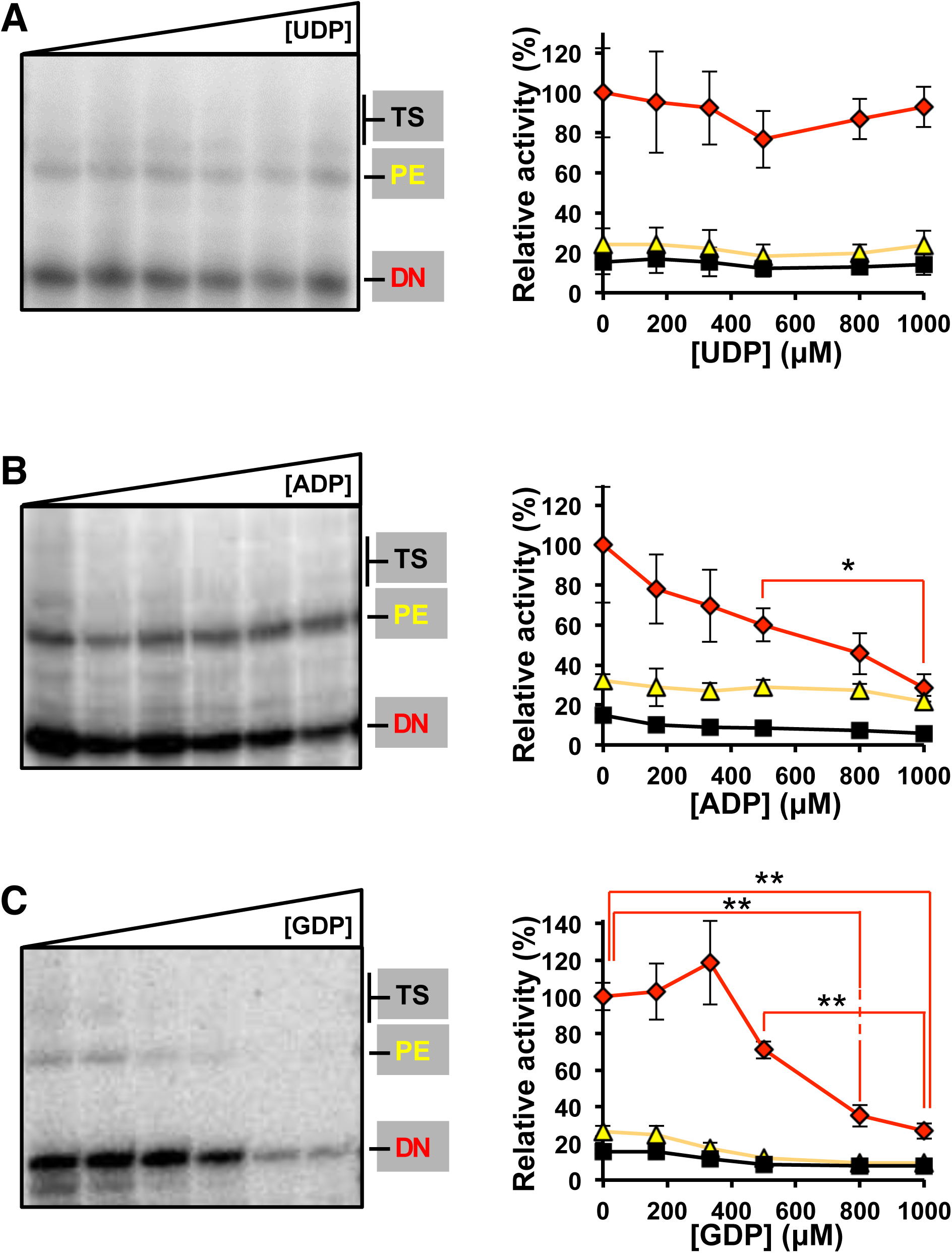
Effect of NDPs on NS5BΔ21 RNA polymerase activity. **(A)** Polyacrylamide gel showing the products for *de novo* (DN), primer extension (PE) and template switching (TS) obtained with HCV NS5BΔ21 at increasing concentrations (0, 166, 333, 500, 800, and 1000 μM) of UDP in the presence of ATP and UTP at a final concentration of 100 μM, GTP at 500 μM, and radiolabeled α^32^P-CTP (left panel). Graphic representation of densitometric values obtained from the electropherogram shown in A (red diamonds, yellow triangles, and black squares correspond to *de novo* (DN), primer extension (PE), and template switching (TS) activities, respectively) (right panel). **(B)** Corresponds to experiments as in **A** but for increasing concentrations of ADP. **(C)** corresponds to experiments as in **A** but for increasing concentrations of GDP. Activities were normalized to their maximum values. Densitometric data represent the mean of at least three independent experiments. Error bars correspond to standard error of the mean. Horizontal lines indicate statistically significant differences (Student’s T-test) between activity values, using the same color code as the activity type. Details of the activity measurements are given in Materials and Methods. Significance (Student’s T-test): ** p < 0.005, * p < 0.05.

### Mutational effects of guanosine

To determine if Gua-related nucleotide pool effects were associated with the mutation repertoire exhibited by HCV during replication in Huh-7.5 cells, the mutant spectrum of the genomic region spanning the last 49 nucleotides of the NS4B gene and the first 490 nucleotides of the NS5A gene, was analyzed using molecular cloning and Sanger sequencing. Following three passages in absence or presence of Gua, the maximum mutation frequency resulted in a significant increase in the HCV populations passaged in the presence of Gua (p<0.0001 and p=0.01 for Gua 500 µM, and Gua 800 µM, respectively; χ^2^ test) (Table 3). A hallmark of virus extinction by lethal mutagenesis is the decrease of specific infectivity (the ratio between viral infectivity and the amount of genomic viral RNA) [7]. Extinction by Gua occurred with a 2.8-fold to 11.8-fold decrease of specific infectivity in the first two passages in the presence of the compared drug, as quantified by infectivity and viral RNA in samples of the cell culture supernatants (Fig 8), suggesting that an increase in polymerase error rate was involved. The most remarkable change was that replication in the presence of Gua increased significantly the number of indels in the heteropolymeric genomic regions of the mutant spectrum (Table 4). Indels in homopolymeric regions ─consisting of at least three successive identical nucleotides─ were not considered because control experiments revealed that they can be amplification artifacts [19]. No indels were detected in the 53 molecular clones derived from the population passaged in the absence of Gua. In sharp contrast, 10 deletions and 2 insertions were present in the 64 molecular clones retrieved from the population passaged in the presence of 500 µM Gua, and 5 deletions in 68 molecular clones from the population passaged in the presence of 800 µM Gua (Table 4). The difference in the number of deletions is highly significant for the populations passaged in the presence of 500 µM and 800 µM Gua (p<0.001; test χ2). The size of the deletions ranged from 1 to 46 nucleotides, some were found in a single clone, others in several clones, and none of the deletions and insertions were in phase; they all generated a premature STOP codon (Table 4 and Fig 9). Therefore, the anti-HCV effect of Gua, exerted via nucleotide-mediated alterations of polymerase activity, is associated with the generation of multiple deletions during HCV replication.

**Figure 8.**
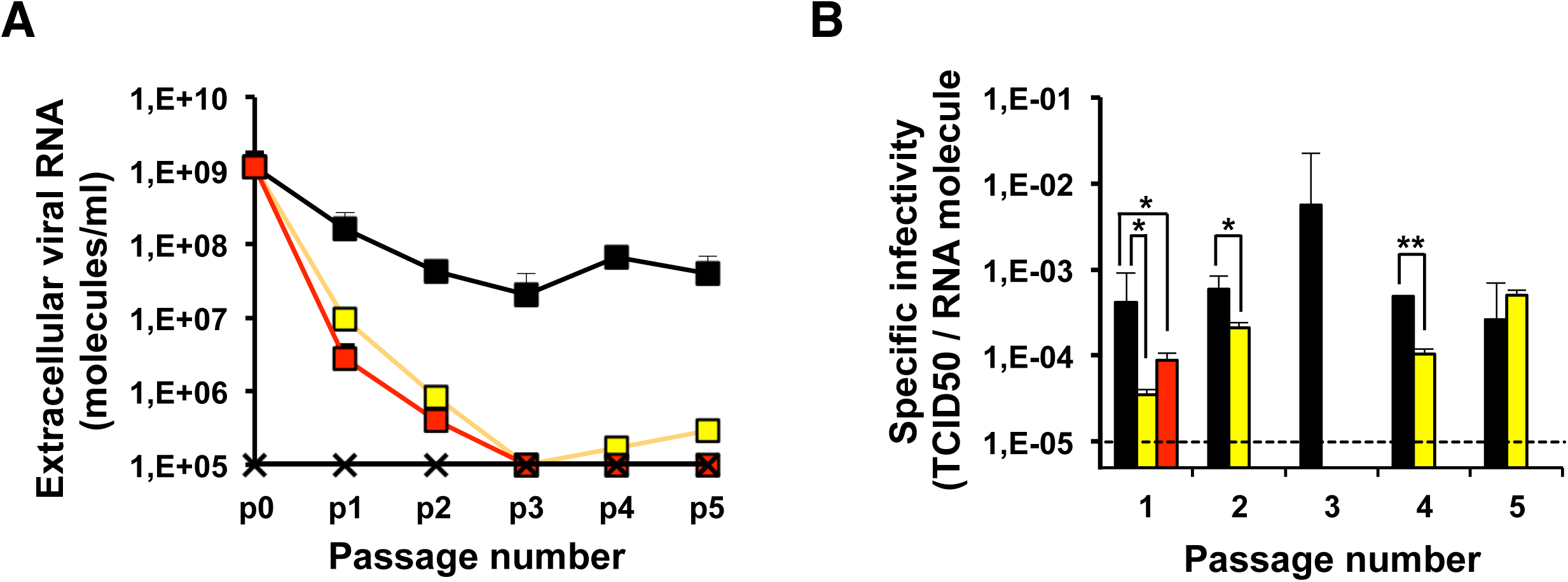
Effect of guanosine on HCV specific infectivity. Huh-7.5 reporter cells were infected with HCVp0 at an initial m.o.i. of 0.05 TCID_50_/cell, in the absence or presence of Gua at the indicated concentrations. HCV GNN infection was used as a negative control. **(A)** Extracellular viral RNA measured by quantitative RT-PCR in different passages. The populations correspond to those of the experiment described in Fig 1 and the values in each passage are the average of the three replicas; standard deviations are given. **(B)** Specific infectivities calculated from the infectivity values of the Fig. 1A and Fig. 1B and the extracellular RNA concentrations indicated in Fig 8A. The horizontal dashed line indicates the limit of detection of viral RNA and specific infectivity. Black, yellow, and red symbols correspond to no drug, Gua 500 μM, and Gua 800 μM, respectively. Values for a HCV lethal mutant GNN (black crosses) are also shown. Details of the procedures are given in Materials and Methods. Significance (Student’s T-test): ** p < 0.005, * p < 0.05.

**Figure 9.**
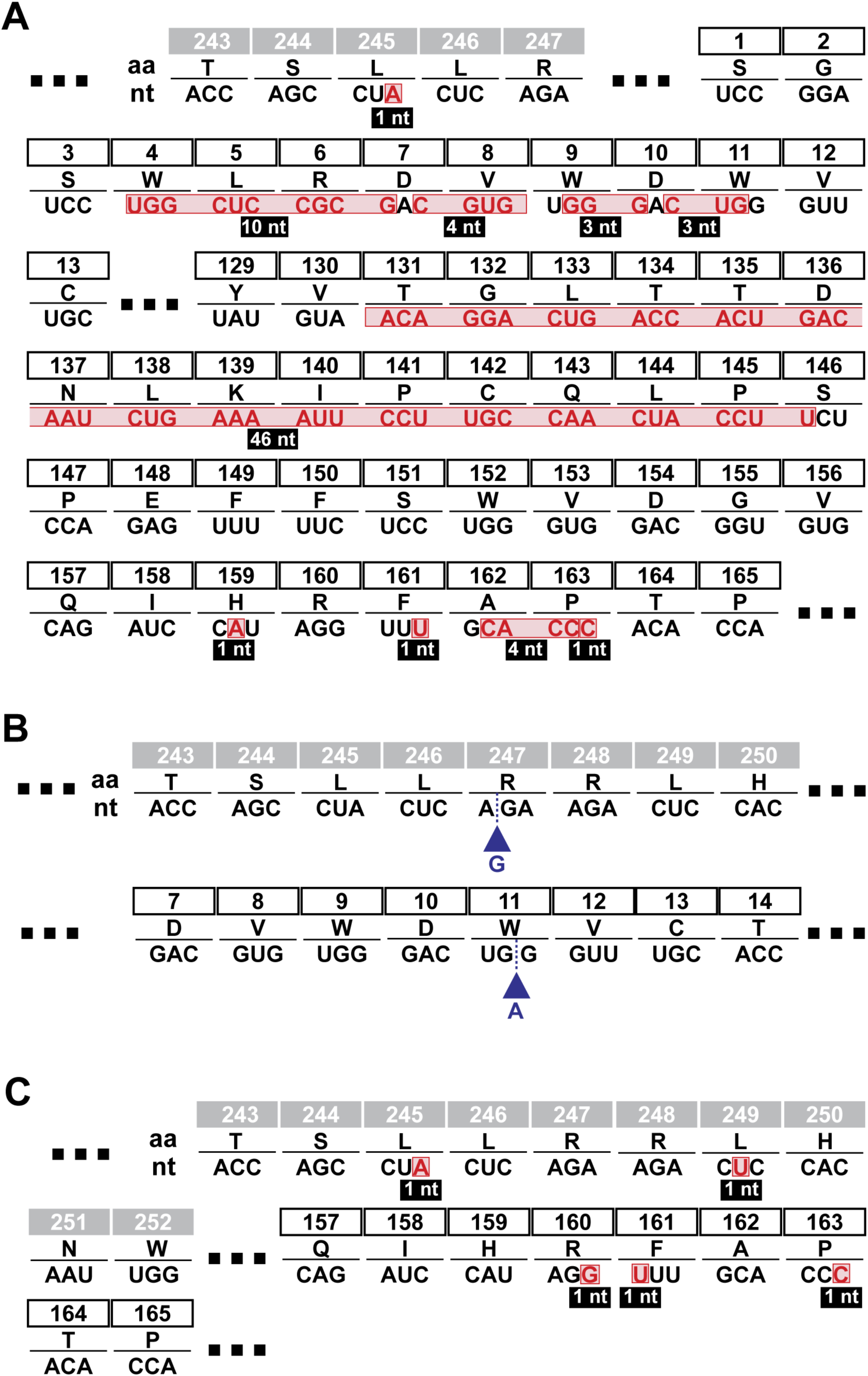
Indels found in the mutant spectrum of HCV p0 passaged in the presence of Gua. The nucleotide sequence of HCV genomic residues 6220 to 6758 was determined for 53 molecular clones derived from the population in absence of Gua, and 132 molecular clones from populations passaged in presence of Gua (data in Table 3). Deduced amino acids (single letter code) are given for residues located at the carboxy-terminal region of NS4B preceding NS5A amino acids. For clarity, only residues around insertions or deletions are shown; three squared points indicate missing amino acids (sequence is that of JFH-1; accession number AB047639). No indels were detected in the population passaged in absence of Gua. (**A**) Deletions in the population passaged in the presence of 500 µM guanosine. Red boxes indicate nucleotides that were deleted in a component of the mutant spectrum, with the deletion size indicated in the filled boxes. (**B**) Insertions in the mutant spectrum of the population passaged in the presence of 500 µM Gua are marked with a blue triangle. (C) Deletions found in the HCV populations passaged in the presence of 800 µM Gua. Procedures for HCV genome sequencing are described in Materials and Methods.

**Table 3.** Mutant spectrum analysis of the hepatitis C virus populations passaged in the absence and presence of guanosine (Gua)

| HCV population<sup>a</sup> | N. of nucleotides analyzed<br>(clones/haplotypes)<sup>b</sup> | N. of different<br>(total) mutations<sup>c</sup> | Mutation frequency |  |
| --- | --- | --- | --- | --- |
|  |  |  | Minimum<sup>d</sup> | Maximum<sup>e</sup> |
| HCV p0, [no drug] p3 | 28,208 (53/21) | 23 (25) | 8.2 x 10<sup>-4</sup> | 8.9 x 10<sup>-4</sup> |
| HCV p0, [Gua 500 µM] p3 | 33,339 (64/34) | 45 (79) | 1.4 x 10<sup>-3</sup> | 2.4 x 10<sup>-3</sup> |
| HCV p0, [Gua 800 µM] p3 | 35,606 (68/30) | 32 (59) | 9.0 x 10<sup>-4</sup> | 1.7 x 10<sup>-3</sup> |

<sup>a</sup>The
<sup>b</sup>The genomic region analyzed by molecular cloning-Sanger sequencing spans residues 6220 to 6758 of the NS4B- and NS5A-coding region; the residue numbering corresponds to the JFH-1 genome (GenBank accession number #AB047639). The values in parenthesis indicate the number of clones analyzed followed by the number of haplotypes (number of different RNA sequences) found in the mutant spectrum.
<sup>c</sup>Number of different and total mutations identified by comparing the sequence of each individual clone with the consensus sequence of the corresponding population.
<sup>d</sup>Data represent the average number of different mutations (counted relative to the consensus sequence of the corresponding population) per nucleotide in the components of the mutant spectrum. The statistical significance of the differences between two populations ( $\chi^2$ test) is the following: HCV p0, [no drug] p3 versus HCV p0, [Gua 500 µM] p3: p = 0.0622; HCV p0, [no drug] p3 versus HCV p0, [Gua 800 µM] p3: p = 0.8257; HCV p0, [Gua 500 µM] p3 versus HCV p0, [Gua 800 µM] p3: p = 0.0977.
°Data represent the average number of total mutations (counted relative to the consensus sequence of the corresponding population) per nucleotide in the components of the mutant spectrum relative to the consensus sequence of the corresponding population. The statistical significance of the differences between two populations (proportion test) is the following: HCV p0, [no drug] p3 versus HCV p0, [Gua 500 µM] p3: $p < 0.0001$ ; HCV p0, [no drug] p3 versus HCV p0, [Gua 800 µM] p3: $p = 0.0107$ ; HCV p0, [Gua 500 µM] p3 versus HCV p0, [Gua 800 µM] p3: $p = 0.0452$

**Table 4.**
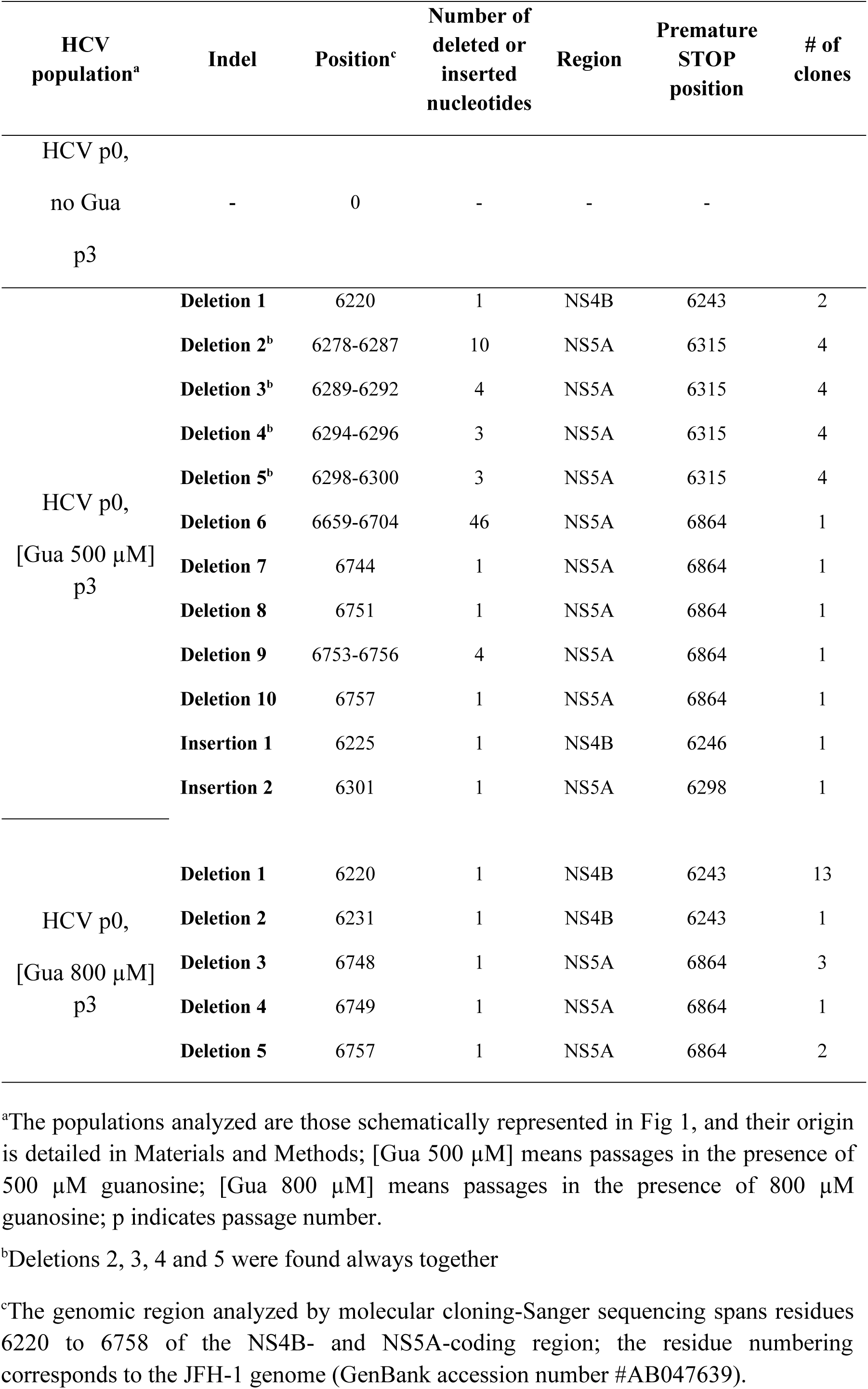
Indels found in the mutant spectra of HCV p0 after 3 passages in the absence and presence of Gua 500 µM and 800 µM.

## Discussion

Nucleoside derivatives are the most important family of drugs targeting viral polymerases, but the antiviral capacity of natural nucleosides has not been described [5]. Interestingly, we observed an inhibitory effect of Gua when it was used in experiments to analyze the impact of mycophenolic acid and ribavirin on HCV progeny production [11]. Here, we document inhibition of HCV replication by Gua in single and serial infections of Huh-7.5 cells that led to loss of infectivity without significant toxicity for the host cells. The antiviral action of Gua was also exerted on high fitness HCV, albeit without loss of infectivity after 5 passages in the presence of Gua, in agreement with the drug resistance phenotype displayed by high fitness HCV (Fig 1) [14–17]. The antiviral effect of Gua was not observed for FMDV in BHK-21 cells or for LCMV and VSV in Huh-7.5 cells (Fig 3).

RNA synthesis by NS5BΔ21 was not significantly affected by Gua concentrations up to 1 mM. Therefore, the inhibition of virus progeny production is unlikely to be the result of direct polymerase inhibition by Gua. This result is consistent with previous work that showed the ability of NS5B to initiate RNA synthesis with this nucleoside [20]. In contrast to Gua, altered intracellular nucleotide concentrations affected the activity of NS5B, in particular an alteration of the *de novo* RNA synthesis by the GTP/GDP and/or ATP/ADP balances (Fig 7). The NS5B protein has an allosteric binding site of GTP and the balance between NDP and NTP might modulate RNA synthesis through this site [21–23].

Some GTP-dependent proteins play a role in the HCV replication cycle. They include proteins involved in virus entry into the cells (i.e HRas), proteins involved in translation (i.e.the eIF5B factor), or in replication (i.e. GBF1) [24–26]. The GTP-dependent Rab18 protein, which is located in the lipid droplets, is involved in capturing proteins such as viral protein NS5A into the replication complex [27]. The observed increase in GTP concentration in Gua-treated cells would not adversely affect the functionality of these proteins.

Some antiviral drugs exert their activity through alterations in the intracellular nucleotide pools [6, 28]. Treatment of Huh-7.5 reporter cells with Gua produced an overall increase of intracellular nucleoside di- and tri-phosphate concentrations (Fig 5). An increase of ATP would be beneficial for virus replication as ATP increments have been associated with the formation of the HCV replication complex [29]. Using previously reported cell volume values for HepG2 cell line (20) (ranging from 2.54 pL to 2.96 pL), the intracellular ATP concentration in Gua-treated Huh7.5-reporter cells would be in the range of 2 mM (Supplementary Table), similar to the previously described values in the HCV replication sites [29].

Intracellular concentration of mono- and di-phosphate nucleotides also modulates metabolic pathways critical for virus replication. For example, HCV proteins NS4B and NS5A inhibit the cellular protein AMP-activated protein kinase (AMPK) [30]. Inhibition of AMPK induces the synthesis of fatty acids and cholesterol that are of vital importance in the HCV replication sites. ADP activates AMPK [31, 32], and the inactivation of AMPK by NS4B and NS5A is ineffective when ADP is increased [30]. As a result, there is no longer accumulation of fatty acids and cholesterol, and viral replication stops. Metformin activates AMPK by increasing AMP and ADP, and this effect has been associated with inhibition of HCV replication. [31,33,34]. The AMPK dissociation constant for ADP is in the range of 1.3-1.5 μM [32]. Therefore, the increase of the ADP concentration from 40 μM to 200 μM (Fig 4 and Supplementary Table) could be preventing, at least in part, inactivation of AMPK. Whether ATP acts only as a substrate or it also exerts some regulatory role needs to be further analyzed.

Defective viral genomes are increasingly recognized as players in virus-host interactions (reviewed in [35]). They are associated with multiple types of genetic lesions ranging from point mutations to large deletions. Deletions can result from polymerase slippage over one or several nucleotides, but the environmental factors that may trigger their occurrence are unknown. Generation of RNA deletions has been documented during *in vitro* replicase copying of viral RNAs [36], and deletions have been observed in many viruses [35,37–40]. Template switching is considered as the primary mechanism of copy-choice recombination of poliovirus in cells [41], and the primary mechanism of poly(rU) RNA synthesis by poliovirus polymerase [42]. The observed increase of indels in the presence of high GTP concentrations may be linked to the enhanced template switching observed with the HCV polymerase *in vitro.* Despite limited information on the origin of HCV defective genomes, there is solid evidence of the implication of defective viral genomes in the interference with replication of their standard infectious counterparts [43], including a contribution to viral extinction by lethal mutagenesis [44, 45]. They also play a role as stimulators of antiviral responses [46], and as mediators of virus attenuation and persistence [40,47,48], among other functions [35].

Defective genomes have been described for HCV, including in-frame deletion mutants. They are present in patient plasma, exosomes and liver biopsies and they may play regulatory roles during viral replication [49–53]. Little is known about the molecular mechanisms of generation of defective genomes despite detailed accounts of their high m.o.i.-dependent selection based on molecular complementation with standard genomes [35]. Our results provide evidence of a mechanism of generation of defective HCV genomes fuelled by nucleoside pool effects on HCV polymerase activity. This is accompanied by a significant reduction of the specific infectivity of the passaged viral pools, demonstrating the increasing presence of non-infectious viral genomes in the supernatants of Gua-treated cells. In addition to unveiling a possible mechanism of generation of defective HCV genomes, our results open the possibility that the alteration of cellular metabolic pathways may be a complementary strategy to the action of antiviral agents to produce reductions in viral load and promote the extinction of HCV.

## Material and methods

### Reagents and plasmids

Nucleosides Ade, Cyt, Gua, and Uri, as well as nucleoside di- and tri-phosphates were purchased from Sigma-Aldrich. Plasmid pNS5BΔ21 encoding the HCV NS5B that lacks the C-terminal 21 hydrophobic amino acids to enhance solubility has been described previously [54]. The resulting expression vector allows the expression of a tagged NS5BΔ21 with six histidine residues at its C terminus to aid in protein purification.

### Cells and viruses

The origin of Huh 7.5, Huh 7-Lunet, Huh-7.5 reporter cell lines and procedures used for cell growth in Dulbecco’s modification of Eagle’s medium (DMEM), have been described [11]. Cell lines were incubated at 37°C and 5% CO2. We used the following viruses in the experiments: HCV p0, obtained from HCVcc [Jc1FLAG2(p7-nsGluc2A)] (genotype 2a) and GNN [GNNFLAG2(p7-nsGluc2A)] (a replication-defective virus with a mutation in the NS5B RNA-dependent RNA polymerase) [11, 55]. Mock-infected cells maintained in parallel with the infected cultures were prepared to control the absence of contaminations; no infectivity in the mock-infected cultures was identified in the experiments.

Trans-encapsidated HCV virions (HCVtcp) were produced by electroporation into packaging cells of a subgenomic, dicistronic HCV replicon bearing a luciferase gene, as previously described [12]. Supernatants of the electroporated cells were titrated to determine the optimal dose rendering detectable luciferase activity at 48 hours post-inoculation. The same subgenomic replicon was used for lipofection experiments, using lipofectamine2000 transfection reagent as previously described [13].

### Production of viral progeny and titration of infectivity

The procedures used to obtain the initial virus HCV p0 and for serial infections of the hepatoma Huh-7.5 reporter cells have been described [14]. Briefly, electroporation of Huh-7 Lunet cells was performed with 10 μg of the transcript of HCVcc (Jc1 or the negative control GNN) (260 volts, 950 μF). Then, electroporated cells were passaged every 3–4 days before cells became confluent; passages were continued until 30 days post-electroporation. Subsequently, the cell culture supernatants were pooled to concentrate the virus 20 times using 10,000 MWCO spin columns (Millipore), and aliquots were stored at −70°C [14]. For titration of HCV infectivity, cell culture supernatants were serially diluted and applied to Huh-7.5 cells. After 3 days post-infection the cell monolayers were washed with PBS, fixed with ice-cold methanol, and stained with anti-NS5A monoclonal antibody 9E10 [14].

### Toxicity test and inhibitory concentration

The CC_50_ was calculated by seeding 96-well plates with Huh-7.5 cells and exposing them to the compound under study during 72 hours. MTT [3-(4,5-dimethylthiazol-2-yl)-2,5-diphenyltetrazolium bromide] was added at 500 μg/ml; after 4 h crystals were dissolved with 100 μl of DMSO and the O.D. measured at 550 nm; 50% cytotoxicity was calculated from quadruplicate determinations [11].

IC_50_ values were calculated relative to the controls without treatment (defined as 100% infectivity) [56]. Determinations were performed in triplicate.

### RNA extraction, cDNA synthesis, PCR amplification, and nucleotide sequencing

Intracellular RNA was obtained from infected cells using the Qiagen RNeasy kit (Qiagen, Valencia, CA, USA). RNA from cell lysates or cell culture supernatants was extracted using the Qiagen QIAamp viral RNA mini kit (Qiagen, Valencia, CA, USA). Reverse transcription (RT) of different HCV genomic regions was performed using avian myeloblastosis virus (AMV) reverse transcriptase (Promega), and subsequent PCR amplification was carried out using AccuScript (Agilent Technologies), with specific primers. Primers for the HCV amplification and the sequencing have been described [11,14,17,19]. Agarose gel electrophoresis was used to analyze the amplification products, using HindIII-digested Φ-29 DNA as a molecular mass standard. In parallel, mixtures without template RNA were reverse transcribed and amplified to monitor the absence of cross-contamination by template nucleic acids. Nucleotide sequences of

HCV RNA were determined on the two strands of the cDNA copies [11, 55]; only mutations detected in the two strands were considered. To analyze the complexity of mutant spectra by molecular cloning and Sanger sequencing, HCV RNA was extracted and subjected to RT-PCR to amplify the NS5A-coding regions, as has been previously described [11]. Amplifications with template preparations diluted 1:10 and 1:100 were performed to ensure that an excess of template in the amplifications was used in the mutant spectrum analysis; the molecular cloning was performed from the undiluted template only when the 1:100-diluted template produced also a DNA band; this procedure avoids complexity biases due to redundant amplifications of the same initial RNA templates [11]. Control analyses to confirm that mutation frequencies were not affected by the basal error rate during amplification have been previously described [57]. Amplified DNA was ligated to the vector pGEM-T (Amersham) and used to transform *Escherichia coli* DH5α; individual colonies were taken for PCR amplification and nucleotide sequencing, as previously described [56].

### NDP and NTP pool analysis

The procedure used has been previously described [11]. Briefly, Huh-7.5 cells (2×10^6^ cells) were washed with PBS and incubated with 900 μl of 0.6 M trichloroacetic acid on ice for 10 min. A precooled mixture of 180 µl of Tri-n-octylamine (Sigma) and 720 µl of Uvasol® (1,1,2-trichlorotrifluoroethane, Sigma) was added to the 900 µl extract, vortexed for 10 s, centrifuged 30 s at 12,000 × g at 4 °C, and stored at −80 °C prior to analysis. One hundred µl samples were applied to a Partisil 10 SAX analytical column (4.6 mm×250 mm) (Whatman) with a Partisil 10 SAX guard cartridge column (4.6×30 mm) (Capital HPLC) using an Alliance 2695 HPLC system connected to a 2996 photodiode array detector (Waters). NDP and NTP were separated at a eluent flow rate of 0.8 ml/min and detected with ultraviolet light at a wavelength of 254 nm. The column was pre-equilibrated with 60 ml of 7 mM NH_4_H_2_PO_4_, pH 3.8 (buffer A). The separation program started with 22.5 min of an isocratic period with buffer A, continued with a linear gradient of 112.5 min to the high concentration buffer 250 mM NH_4_H_2_PO_4_, 500 mM KCl, pH 4.5 (buffer B) and ended with an isocratic period of 37.5 min with buffer B. A processing method was done using the Waters Empower™ Chromatography Data Software. To this end, 50 µl of 20 pmol/µl UTP, CTP, ATP and GTP (Jena Bioscience), were separated prior to sample analysis. The HPLC analysis did not separate rNTPs from dNTPs, or rNDPs from dNDPs. However, since the absolute concentration of rNTPs and rNDPs is several orders of magnitude greater than that of dNTPs dNDPs, we consider the value obtained as the concentration of rNTPs and rNDPs. Determinations were carried out with two independent biological samples, each one in triplicate for NDPs and NTPs. The amount of each nucleoside in cell extracts was normalized relative to the number of cells.

### Quantification of HCV RNA

Real time quantitative RT-PCR was performed with the Light Cycler RNA Master SYBR Green I kit (Roche), following the manufacturer’s instructions, as previously described [14]. The 5′-UTR non-coding region of the HCV genome was amplified using as primers oligonucleotide HCV-5UTR-F2 (5′- TGAGGAACTACTGTCTTCACGCAGAAAG; sense orientation; the 5′ nucleotide corresponds to HCV genomic residue 47), and oligonucleotide HCV-5UTR-R2 (5′-TGCTCATGGTGCACGGTCTACGAG; antisense orientation; the 5′ nucleotide corresponds to HCV genomic residue 347). Quantification was relative to a standard curve obtained with known amounts of HCV RNA, obtained by *in vitro* transcription of plasmid GNN DNA [55]. Reaction specificity was monitored by determining the denaturation curve of the amplified DNAs. Mixture without template RNA and RNA from mock-infected cells were run in parallel to ascertain absence of contamination with undesired templates.

### NS5BΔ21 polymerase expression and purification

NS5B from strain pJ4-HC with a deletion of 21 aa at the C-terminal end (NS5B**Δ**21) was obtained as previously described [54, 58]. This truncated protein displays polymerase activities that were not distinguished from those of the full-length enzyme [59]. Briefly, NS5B**Δ**21 was overexpressed in BL21DE3 Rosetta bacteria by IPTG induction and purified by affinity chromatography in a Ni-NTA column. Aliquots of the purest and most concentrated protein samples were adjusted to 50% glycerol and stored at -80**°**C until use. All purification processes were monitored by SDS-PAGE and Coomassie brilliant blue staining. Protein was quantified by SDS-PAGE gel imaging and protein determination using the Bradford assay.

### *In vitro* RdRP replication assays

RNA polymerase assays were carried out using two different RNA substrates, the symmetric substrate LE-19 (sequence 5’ UGUUAUAAUAAUUGUAUAC 3’), which is capable of primer-extension (PE), *de novo* initiation (DN), and template switching (TS) [54, 58], and an RNA fragment encompassing HCV E1/E2 region (570 nt) [18]. Except when indicated otherwise, template RNA was pre-incubated for 15 minutes in a reaction mixture containing 20 mM MOPS, pH 7.3, 35 mM NaCl, 5 mM MnCl2, 100 nM NS5B and GTP at the indicated concentration for each experiment. Reactions were started by adding 1 µCi of α[^32^P]CTP (3000 Ci mmol, PerkinElmer Life Sciences) and nucleoside-triphosphate as indicated in each experiment. When appropriate, reactions were performed in the presence of increasing concentrations of nucleotide diphosphates. Reactions were carried out in a final volume of 10 µl, at room temperature for 30 minutes, and stopped using EDTA/formamide loading buffer. E1/E2 products were resolved using 1% agarose gel electrophoresis. Agarose gels were dried in an electrophoresis gel dryer (BioRad). LE19 products were resolved using denaturing polyacrylamide (23% PAA, 7 M urea) gel electrophoresis. Gels were exposed to phosphorimager screens and scanned with a Typhoon 9600 phosphorimager (Molecular Dynamics). Sample quantification was performed from parallel experiments. Band volume values were obtained by using the ImageQuant software provided with the apparatus (GE Healthcare).

### Statistical analyses

The statistical significance of differences between mutation frequencies was evaluated by the chi-square test. Statistical comparisons among groups were performed with Student’s T-tests. Unless indicated otherwise, the statistical significance is indicated by asterisks: * p<0.05; ** p<0.005; *** p<0.0005.

## Acknowledgements.

C.P. is supported by the Miguel Servet program of the Instituto de Salud Carlos III (CP14/00121) cofinanced by the European Regional Development Fund (ERDF). CIBERehd (Centro de Investigación en Red de Enfermedades Hepáticas y Digestivas) is funded by Instituto de Salud Carlos III. The work at CBMSO was supported by grants SAF2014-52400-R from Ministerio de Economía y Competitividad (MINECO), SAF2017-87846-R, BFU2017-91384-EXP from Ministerio de Ciencia, Innovación y Universidades (MICIU), PI18/00210 from Instituto de Salud Carlos III, S2013/ABI-2906, (PLATESA from Comunidad de Madrid/FEDER) and S2018/BAA-4370 (PLATESA2 from Comunidad de Madrid/FEDER). The team at CBMSO belongs to the Global Virus Network (GVN). The work in Albacete was supported by grants SAF2016-80451-P, EQC2018-004420-P, and EQC2018-004631-P from MICIU, and by Plan Propio of the University of Castilla-La Mancha. The work in Malaga was supported by Plan Propio of the University of Málaga. Institutional grants from the Fundación Ramón Areces and Banco Santander to the CBMSO are also acknowledged. Piet de Groot is acknowledged for critical reading the manuscript.

