## Supplemental Table for "Guanosine inhibits hepatitis C virus replication and increases indel frequencies, associated with altered intracellular nucleotide pools"

**Supplementary Table.** Intracellular concentration range (in mM) of nucleoside di- and triphosphates after treatment with Gua 500 and 800 µM.

| [mM]^a^ | No Drug | Gua 500 μM | Gua 800 μM |
| --- | --- | --- | --- |
| UTP | 0.2-0.3 | 0.2-0.3 | 0.4-0.5 |
| CTP | 0.1 | 0.1 | 0.2-0.3 |
| ATP | 0.8-0.9 | 1.0-1.2 | 1.8-2.0 |
| GTP | 0.2 | 0.2 | 0.3-0.4 |
| UDP | 0.1 | 0.3-0.4 | 0.3 |
| CDP | 0.01 | 0.02 | 0.01 |
| ADP | 0.04 | 0.2 | 0.1 |
| GDP | 0.06-0.07 | 0.13-0.15 | 0.13-0.15 |

^a^The concentration range was calculated using cell volume previously reported for the cell line HepG2 [60] and the data from Fig 4.
